# Neural Correlates of Perceptual Grouping in Brain

**DOI:** 10.1101/2025.08.12.669980

**Authors:** Shefali Gupta, Tapan Kumar Gandhi

## Abstract

Humans are proficient in detecting even subtle regularities in their sensory inputs. In the specific context of vision, one manifestation of this ability is the perception of structures embedded in noisy backgrounds. Here we use electrophysiology to explore the neural correlates of this proficiency. Observers underwent EEG recordings while being presented with random-dot displays. A few dots in some of the displays were arranged in a line. The results reveal that the two kinds of displays, those with and without the embedded lines, though identical in low-level attributes such as mean luminance and spatial frequency distribution, elicit markedly different neural responses. We find that the first divergence in the ERPs corresponding to these two stimulus classes occurs around 100 milliseconds. These differences are evident in occipital sensors suggesting that the perceptual distinction between structured and non-structured stimuli is accomplished in the very early cortical stages of visual processing. The short latency and early locus of the ERP divergence point to a largely feedforward computation underlying this feat of perceptual organization. The functional connectivity for difference between the two events also supports the ERP findings and thus, exhibits relation between synchrony time-course and ERP dynamics.

## Introduction

Consider the two panels in figure 1A. Both are similar in their aggregate low-level statistics, such as mean luminance and spatial frequency content. But perceptually they appear very distinct; one has elements organized into a structure, here a line, while the other lacks such organization. Our brain is proficient at detecting such organization in complex input arrays. This is perhaps an important precursor to our well-honed ability to spot objects in our surroundings, even in difficult circumstances such as camouflage [1]- [3]. The phenomenology and principles of perceptual organization have a rich history of study [4]- [8], most prominently in the Gestalt tradition. While there is incontrovertible evidence that perception necessarily involves the detection, or even imposition, of organization on input signals, there is less clarity on the neural substrates underlying these processes [9]. This is the issue we address here.

**Figure 1.**
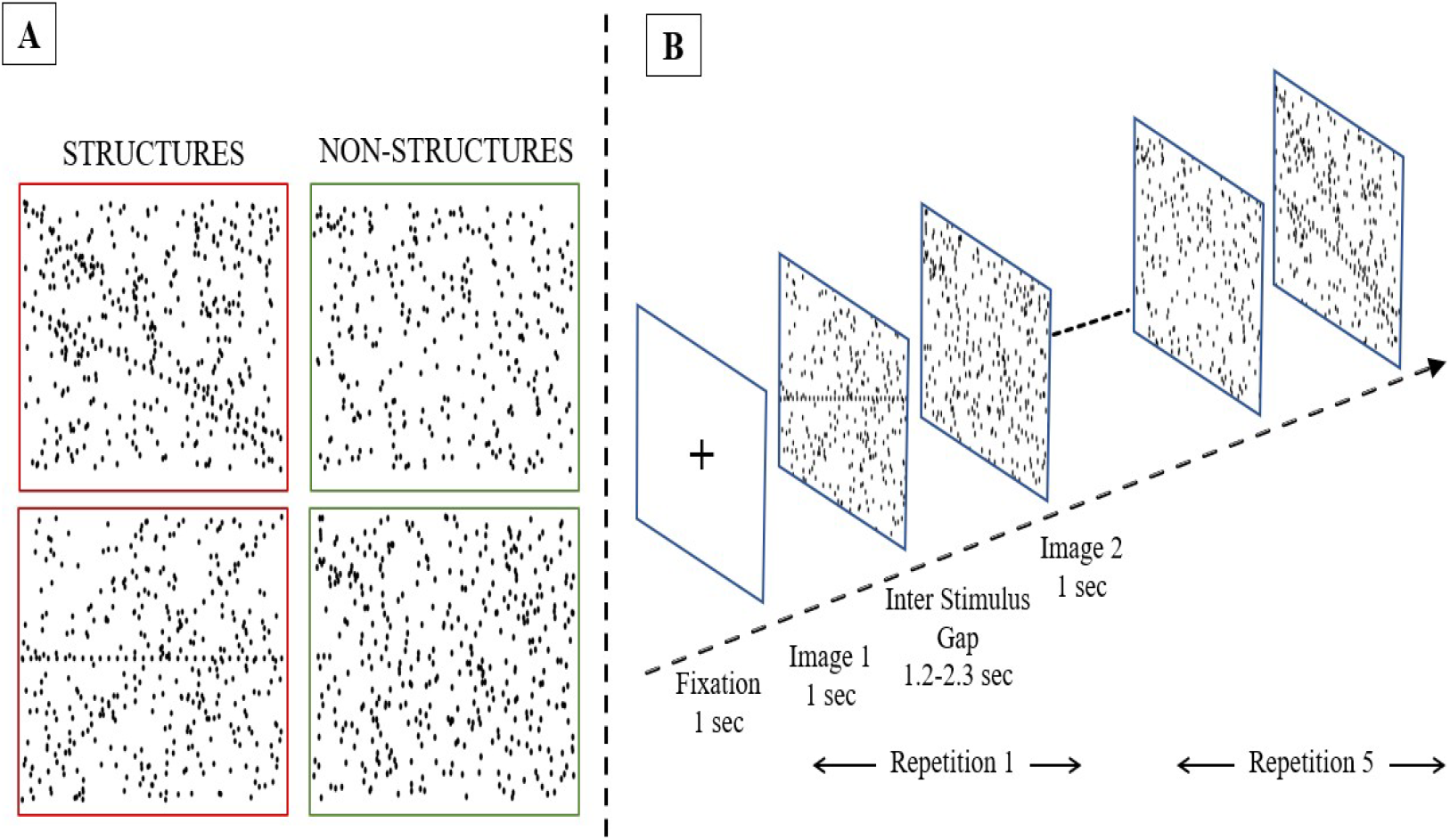
Stimuli Design Overview (A) Examples of stimuli i.e. ‘Structure’ and ‘Non-Structure’; the image displays random dots with some appearing to create perceptual line segments referred to as ‘Structures’ while the other image solely consists of random dots, referred to as ‘Non-Structures’ (B) Outlook of stimulus presentation: Each image was displayed on the screen for 1000ms, followed by an inter-stimulus interval ranging from 1200ms to 2300ms between successive images. In total, 200 stimuli, comprising 100 structure images and 100 non-structure images, were presented randomly to the participants.

We seek to determine whether we can discern differences in non-invasively recorded human brain activity depending on whether the presented stimulus has organized structures or not. Furthermore, if we do find such neural signatures, we would like to use their measured latency and location to make tentative inferences about the nature of neural processing underlying the observed differences. Towards this end, we will adopt the approach of Thorpe and colleagues who used speed of processing as an indicator of whether the associated neural computations were largely feedforward in nature or might also involve feedback loops.

Past work on perceptual organization has focused more on the phenomenology and heuristics that appear to be at work [11]-[12]. Less is known about the neural basis of perceptual grouping [13]-[18]. It is believed that the object to be perceived is extracted from the highly cluttered real-world scenes [19], followed by figure-ground organization [20]-[21]. The components combined to form a shape are coded by neurons in primary visual cortex [22][23].

Various imaging modalities have been employed in previous studies to explore perceptual organization. These include electroencephalography (EEG) [24]-[26], magnetoencephalography (MEG), and functional MRI [27]-[30]. Of particular interest to us is EEG, since that is the technique, we employ in this work. Past studies have typically used conventional ERP analyses, which try to determine when after stimulus presentation a distinct response becomes evident. (A prototypical example is provided by the N170 component. A negative deflection is evident in some sensors over the right temporal lobe about 170 ms following the presentation of a face image). Such analyses have indicated the presence of contour driven ERP components a few hundred milliseconds after stimulus presentation. For example, in a contour-closure determination task [31]-[32], an ERP component with a peak around 290 ms was observed.

Since one of our key goals in this study is to determine how quickly any evidence of perceptual organization associated responses are apparent, we opt for a different style of EEG analyses. These have been used fruitfully by Thorpe and colleagues [10]. They measured the first point of divergence or differential response across stimulus classes. The point of divergence can justifiably be considered evidence of stimulus class discrimination by the brain [33]- [37]. An early divergence in space and time is indicative of feed-forward processing.

With this background, the main objective of the present work is to determine whether differential activity is evoked by structured and non-structured stimuli. Topographical maps showing the dynamics of brain activity can provide clues regarding the feedforward or feedback processing.

The Brain network can be considered as a collection of modules [38] interlinked to each other, so as to carry out different functions [39]. There must be some synchronization between these modules to perceive the object as a whole [40]. Reduction in this synchronization between the modules can cause psychiatric and neurological disorders [41]. One of the coordinating mechanisms found is the synchronization of neuronal activity by phase locking of self-generated network oscillations [42]-[43] known as Phase-locking value (PLV). PLV found application in various neuroscience studies [44]-[45] to quantify the level of PS (Phase Synchronization). Other measures such as phase synchronization index (PSI), correlation coefficient, coherence, mutual information, and partial directed coherence also exists [46]-[50]. In present study, we also tried to find whether the synchrony and the brain response have some relationship for perceptual grouping.

In the present work, we have attempted to provide a deep insight to the concept of perceptual grouping in terms of neural processing. An ERP study is conducted to evaluate the activation of brain response for basic structures (grouping perception) as compared to random dots using EEG. The point of divergence is determined to find the reaction time of brain for perceptual grouping supported by the behavioral study. The functional connectivity analysis of the difference between events is performed over time-length of stimuli presentation to find the PLV-ERP relation dynamics. Topographical Maps are carried out over time-length of stimuli to judge the flow of information during the process of perception formation in brain. Furthermore, the behavioral study signifies how faster the perception formation process can take place in the brain.

## Methods

### Participants

The study was performed with 15 subjects (13 males, 2 females). All the participants were aged between 20 and 30 years. None of them had any neurological diagnoses. The study was approved by the Indian Institute of Technology (IIT) Ethical Clearance Committee review board. Informed consent was obtained from all participants.

### EEG Experiment Design

Figure 1A shows our experimental stimuli, and figure 1B schematizes the timeline of stimulus presentation. Stimuli were of two kinds: structured and non-structured. Both contained 300 randomly positioned black dots on a white background. However, in the structured class, a subset of the dots was organized into two-line segments. Each stimulus was presented for 1 second. The inter-stimulus interval i.e., ISI ranged between 1200 ms to 2300 ms. During ISI, a fixation cross was presented at the center of the screen. A total of 200 stimuli (100 structured and 100 non-structured) were presented in random order. We used the Psychopy software in conjunction with Brain Vision Recorder software to record the EEG signals time locked to stimulus presentation.

### Behavioral Study

The behavioral study was conducted using the same stimuli as in the EEG experiment. The only change in the protocol was that during the ISI, participants were instructed to press ‘1’ if they saw structure (i.e., dots grouping to form two straight lines) and ‘2’ if they did not perceive any lines. The response of participants was recorded along with the reaction time for each presented stimulus.

### EEG data acquisition

The raw EEG data corresponding to stimuli presented was recorded at the Neuro-Computing Lab, IIT Delhi using ActiChamp 64 Channel Brain Vision Products device. Figure 2 shows the EEG acquisition apparatus setup. This comprised two computer systems which were synchronized with each other so that the data were recorded with the trigger code of stimuli presentation. The stimuli were presented to the subject on the PC using Psychopy software while the other PC recorded EEG signals using Brain Vision Recorder software after passing through the amplifier.

**Figure 2.**
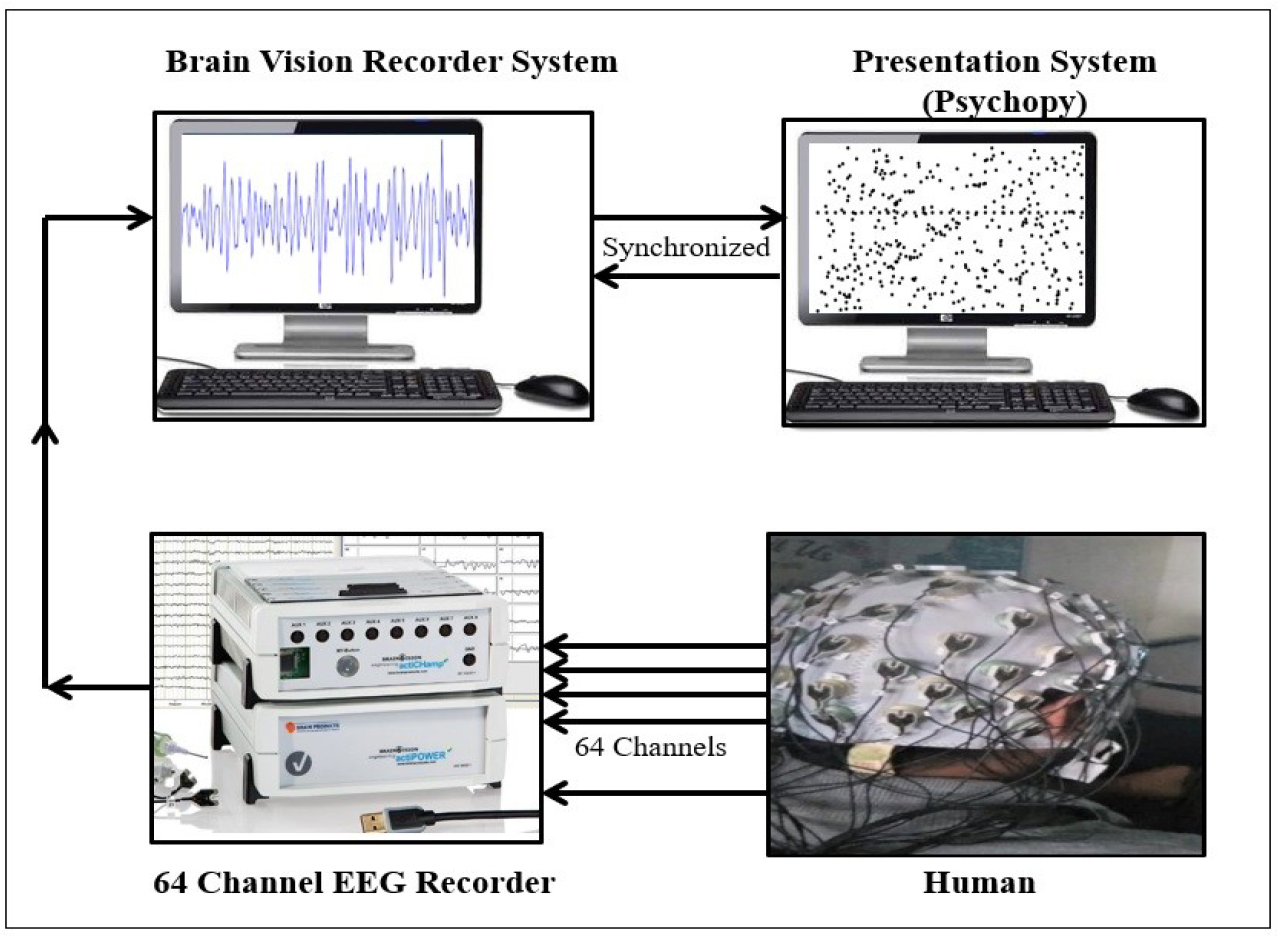
EEG Acquisition Apparatus: The EEG acquisition apparatus setup consists of two computer systems, viz., the recorder system and the presentation system, which are synchronized with each other. The amplifier in an EEG device amplifies the electrical signals detected by the electrodes positioned on the scalp of human participants. The recorder captures and stores the amplified signals received from the brain.

Subject preparation for an EEG session involved head circumference measurement, placement of the appropriately sized electrode cap on the participant’s head, and application of conducting gel to each electrode before the experiment to ensure low-impedance contacts. It took approximately 30-40 minutes to prepare a given subject.

### Preprocessing

The raw EEG data was bandpass filtered with cutoff frequencies 0.5 Hz-45 Hz. The average re-referencing was used to avoid the task related activity to be centered round a particular electrode. Following that each signal was divided into segments of interest for the events specified with the help of triggers send during acquisition. Each epoch is a time-window of length of 1400 ms i.e., from -200 ms before the stimuli onset to 1200 ms after the stimuli onset, extracted from continuous EEG signal. The baseline before the stimuli begins is removed further. Wavelet decomposition is also used for high frequency noise removal.

### ERP analysis

The Event Related Potential (ERP) was evaluated in frontal and occipital region of the brain. The Electrodes showing significant difference between the two events ERPs in frontal and occipital region were determined. The responses of frontal electrodes namely FP1, Fp2, AF7, AF3, AF4, AF8, F5, F6 were averaged. It is found that the Frontal region ERP incorporates three major ERP components namely P200, N400, and a LPC (late positive component) around 800 ms. P200 wave refers to the positive deflection peaking around 150-275 ms after the stimulus onset and is responsible for stimulus classification [51]. N400 wave, also known as FN400 is a negative-going wave that peaks around 300-450 ms post-stimulus during recognition tasks [52]. FN400 indexes repetition-related process and indicates potential for conceptual integration [53]. LPC generally elicited around 600-1000 ms after the stimulus presentation is a positive deflection and is related to more attention or feature-by-feature comparison of stimuli [54]. The resulting Frontal ERP is checked for significant amplitude difference between Structure and Non-Structure events for all its components viz. P200, FN400 and LPC using one-way ANOVA analysis.

The electrodes that showed significant difference between the events at occipital region are-PO7, PO8, PO3, PO4, O1, O2, Oz. The response of these electrodes was averaged to calculate the occipital region ERP. It is observed that ERP at occipital region incorporates four major ERP components namely P100, N200, P300, and a LPC (late positive component) around 800 ms. P100 wave refers to the positive deflection peaking around 90-150 ms after the stimulus onset and is sensitive to visual cognitive processing [55]. N200 wave is a negative-going wave that peaks around 180 ms post-stimulus and is driven either for highly unfamiliar stimuli or stimuli with complex objects [56]- [57]. P300 wave generally elicited 300 ms after the stimulus presentation is a positive deflection and indicates discrimination of one event from another [57]. The resulting Occipital ERP is checked for significant amplitude difference between Structure and Non-Structure events for all its components viz. P100, N200, P300 and LPC using one-way ANOVA analysis.

ERP was also calculated over primary visual cortex (for parieto-occipital region) electrodes. The difference wave showed first negative peak around 70ms after stimulus onset [58]. The other ERP components were same as in occipital ERP.

### Connectivity Analysis

The functional connectivity related to both events was calculated for all 64 electrodes over the time-length of stimuli. The concept of Phase Synchronization (PS) is used to measure the connectivity between electrodes on the scalp. The population of neurons that are phase-locked with each other are said to be wired together and it is believed that the neurons that are wired together, will fire together [59]. Thus, the Phase Locking Value (PLV) reflects the existence of some functional wiring or connectivity between the electrodes. PLV uses Hilbert transform to calculate the instantaneous phase of each signal. The magnitude of phase difference between two electrodes denotes the functional connectivity between them [60]. The phase locking value (PLV) is given as:

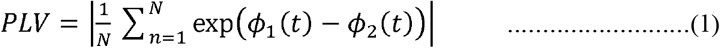

where N is number of trials, *ϕ*_1_(*t*) and *ϕ*_2_(*t*) represents the instantaneous phase of the first and second electrode of the pair respectively. The PLV value ranges between 0 and 1. The high PLV value signifies high functional connectivity between the brain regions. To visualize the PLV values between electrodes, the connectivity matrix and connectivity map is illustrated. For validating the results, the obtained PLV values are compared with the PLV values computed for surrogate data. The surrogate data is generated by randomizing the phase of the original signals [60]. The results were considered only if the difference between surrogate data PLVs and original data PLVs is significant (*p <* 0.05).

## Results

### Behavioral Results

The average of correct responses was 92% while two subjects achieved 97% correct responses. The distribution of reaction time of 15 subjects participated in the experiment over target trials is shown in Figure 3 (A). The median reaction time for ‘structure’ event trials was 450 ms however between subjects it varied from 407 ms to 924 ms. This difference in reaction time between subjects may be because of limited instructions to the participants before experiment and the novelty of the stimuli. The trade-off between reaction time and accuracy for each subject is shown in Figure 3 (B). The point of divergence for ERP difference and the behavioral reaction time showed no correlation between them.

**Figure 3.**
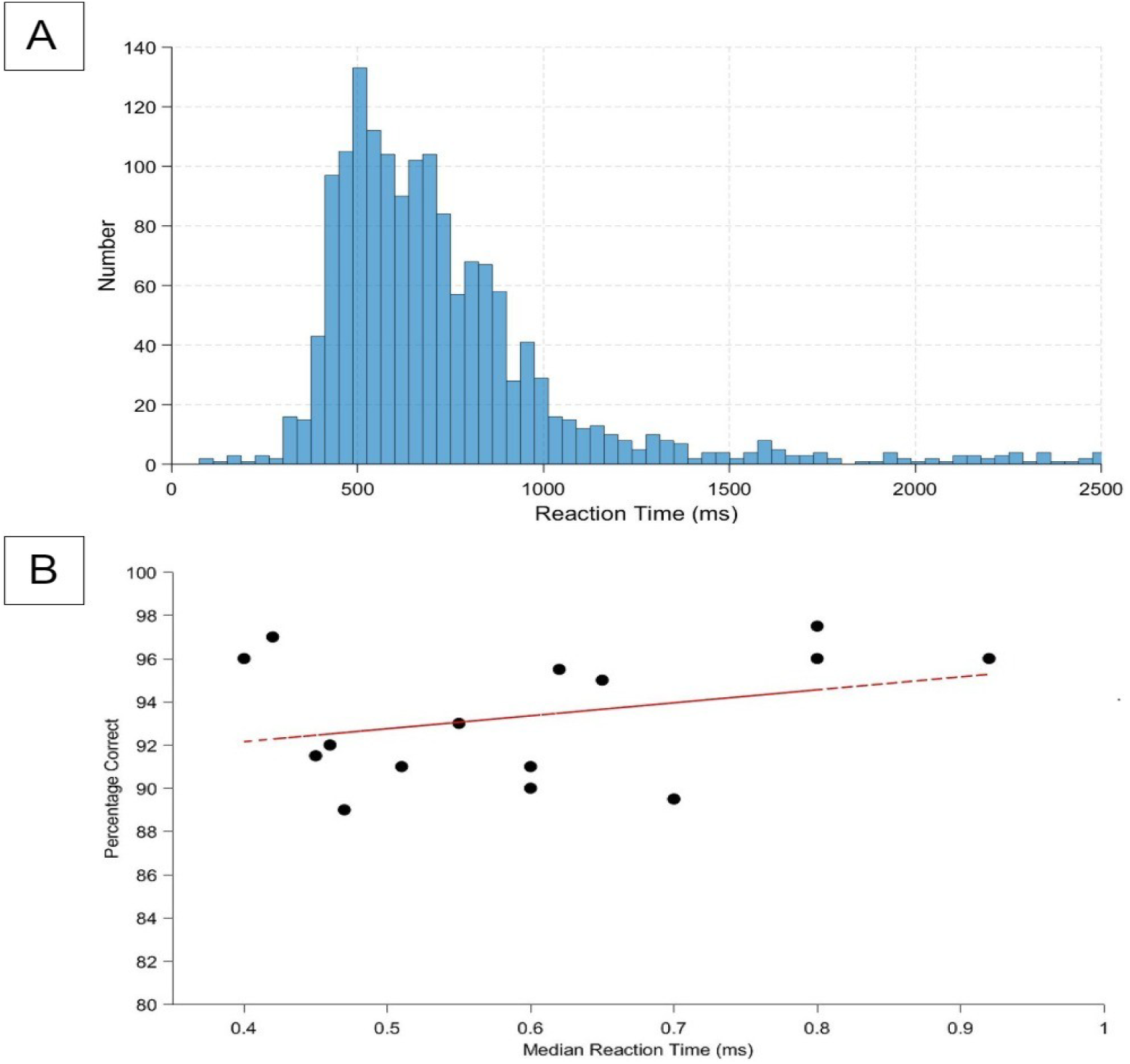
Behavioral Study measures (A) represent distribution of reaction time over number of target trials for all the subjects participated in study (B) represents median reaction time and accuracy trade-off for each subject (dashed line indicates linear regression)

### ERP Results

The Frontal region ERP is shown in Figure 4(A). The Statistical analysis [61]-[62] is done for determining the point of divergence of difference between the events. The point of divergence for frontal region ERP occurs around 100 ms. The first significant point of divergence was observed for electrode F6 at 93 ms. All twelve frontal electrodes reach significance by 135ms (1.2 < t < 3.44). According to T-Score Analysis, the level of significance reaches a peak around 307 ms where the mean t-score for these electrodes was 12.5. The Occipital region ERP is shown in Figure 4 (B). The potential for two events diverges sharply for occipital region around 60 ms after stimulus onset. The Statistical analysis [61]-[62] is done for determining the point of divergence of difference between the events. The first significant divergence point was obtained for electrode O1 at 54ms. All seven occipital electrodes reach significance by 61ms (1.78 < t < 3.55). According to T-Score Analysis, the level of significance reaches a peak around 330 ms where the mean t-score for these electrodes was 28.3.

**Figure 4.**
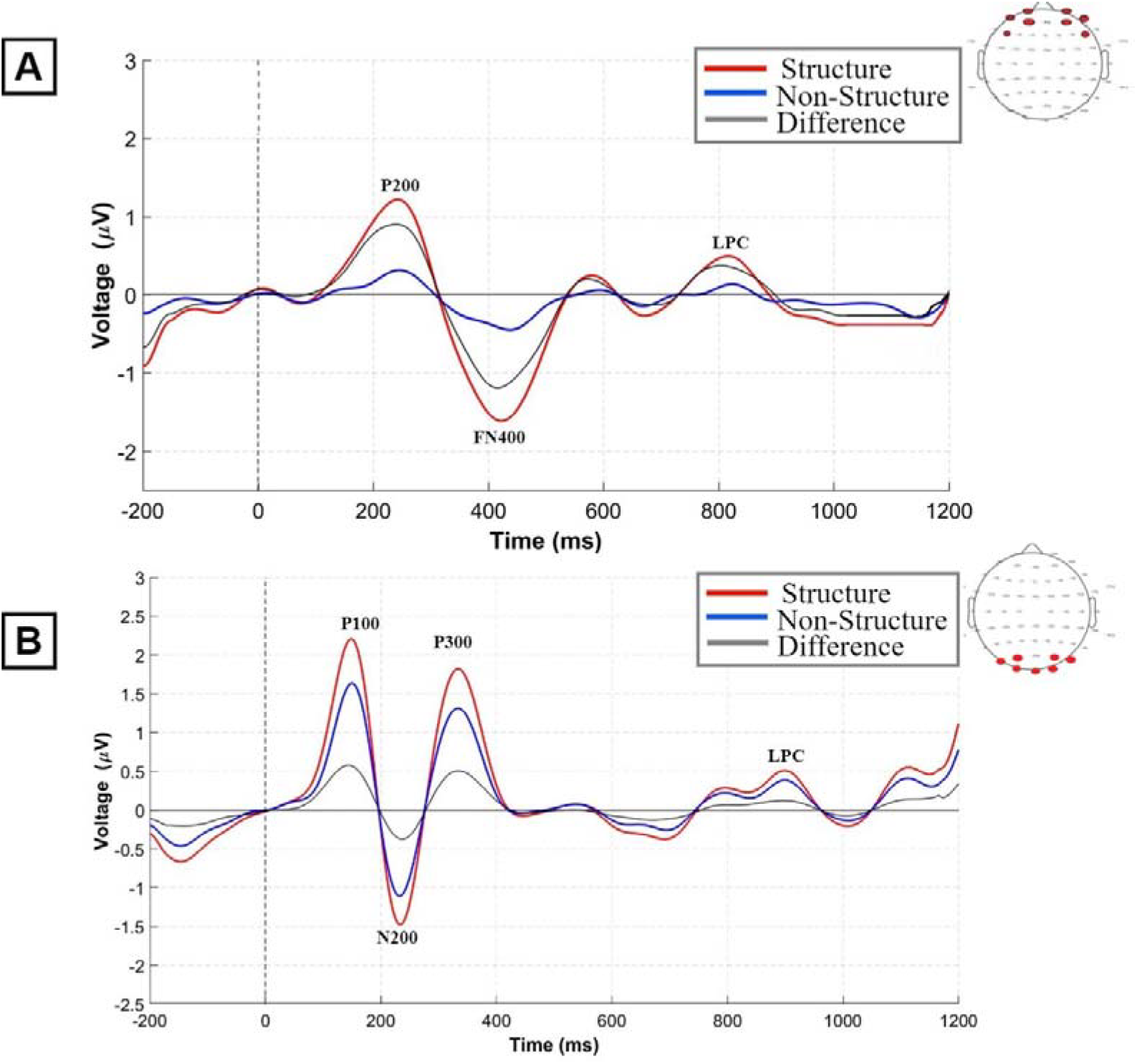
(A) ERP at the Frontal region averaged for electrodes Fp1, Fp2, AF7, AF3, AF4, AF8, F5, and F6 (depicted in the scalp map in the top right corner) following the presentation of Structure-NonStructure stimulus. ERP components P200, FN400, and LPC exhibit larger amplitudes in response to structures (red line) compared to non-structures (blue line). The black line shows the difference between the two events (B) ERP at the Occipital region averaged for electrodes PO7, PO8, PO3, PO4, O1, O2, and Oz (represented in the scalp map at the top right corner) following the presentation of Structure-NonStructure stimulus. ERP components P100, N200, P300, and LPC exhibit larger amplitudes in response to structures (red line) compared to non-structures (blue line). The black line shows the difference between the two events.

### Grand ERP Results

The Grand ERP for Frontal region electrodes for 15 subjects is shown in Figure 6(A). The Statistical analysis [61]-[62] is done to evaluate the initial point of differential response across 15 subjects. The point of divergence for frontal region GERP occur around 90 ms (electrode Fp1), shown in Figure 6(A). All twelve frontal electrodes showed consistent difference from 85ms (1.3 < t < 3.12). The difference curves for all 15 subjects is shown in Figure 6(C). The significance reaches a peak at 335ms where the mean t-score was 22.07, as illustrated in Figure 6(E).

**Figure 5.**
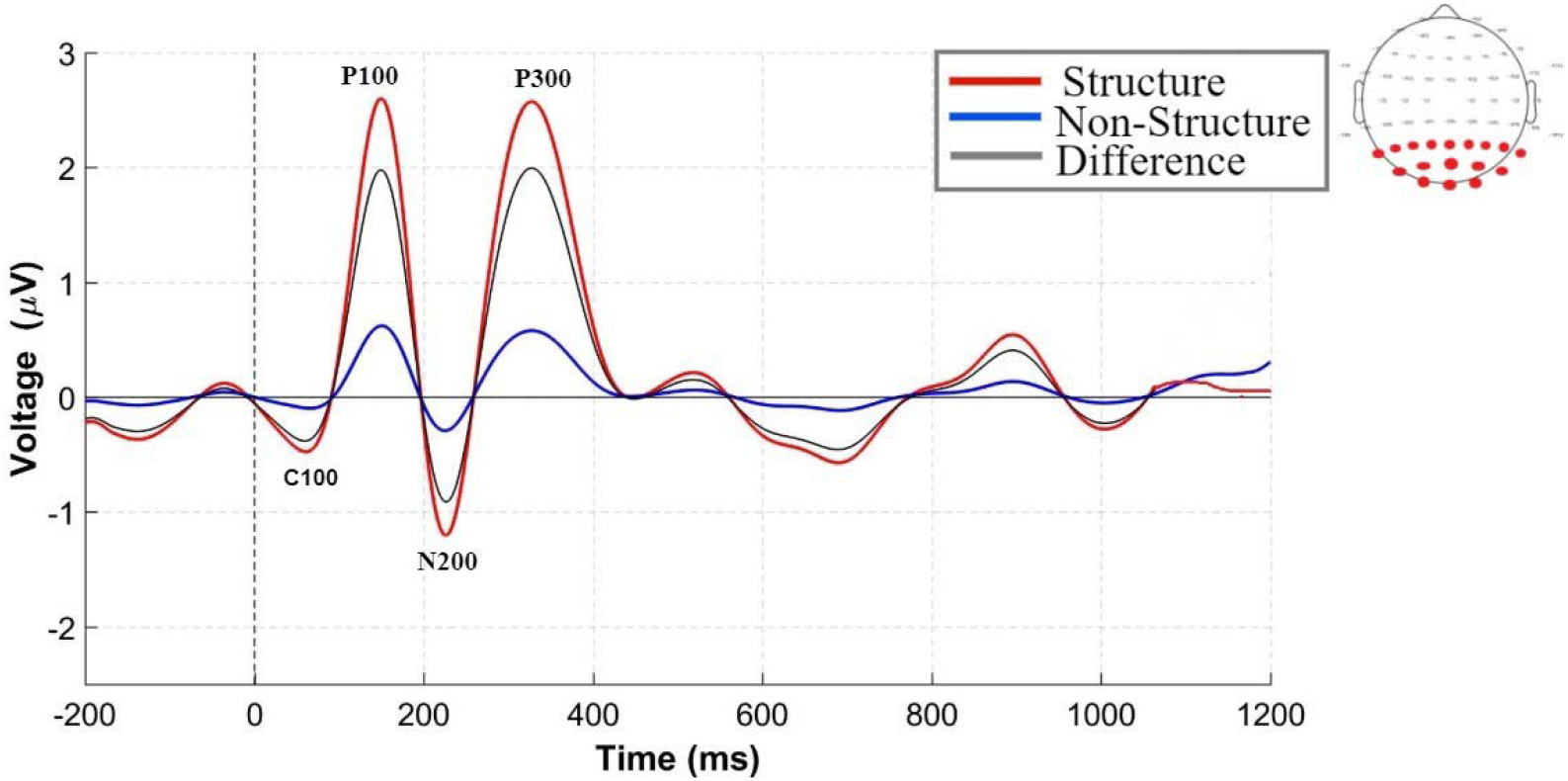
ERP at the V1 region averaged for electrodes P7, P5, P3, P1, Pz, P2, P4, P6, P8, PO7, PO8, POz, PO3, PO4, O1, Oz, and O2 (represented in the scalp map at the right corner) following the presentation of Structure-NonStructure stimulus. ERP components P100, N200, and P300 exhibit larger amplitudes in response to structures (red line) compared to non-structures (blue line). The black line shows the difference between the two events.

**Figure 6.**
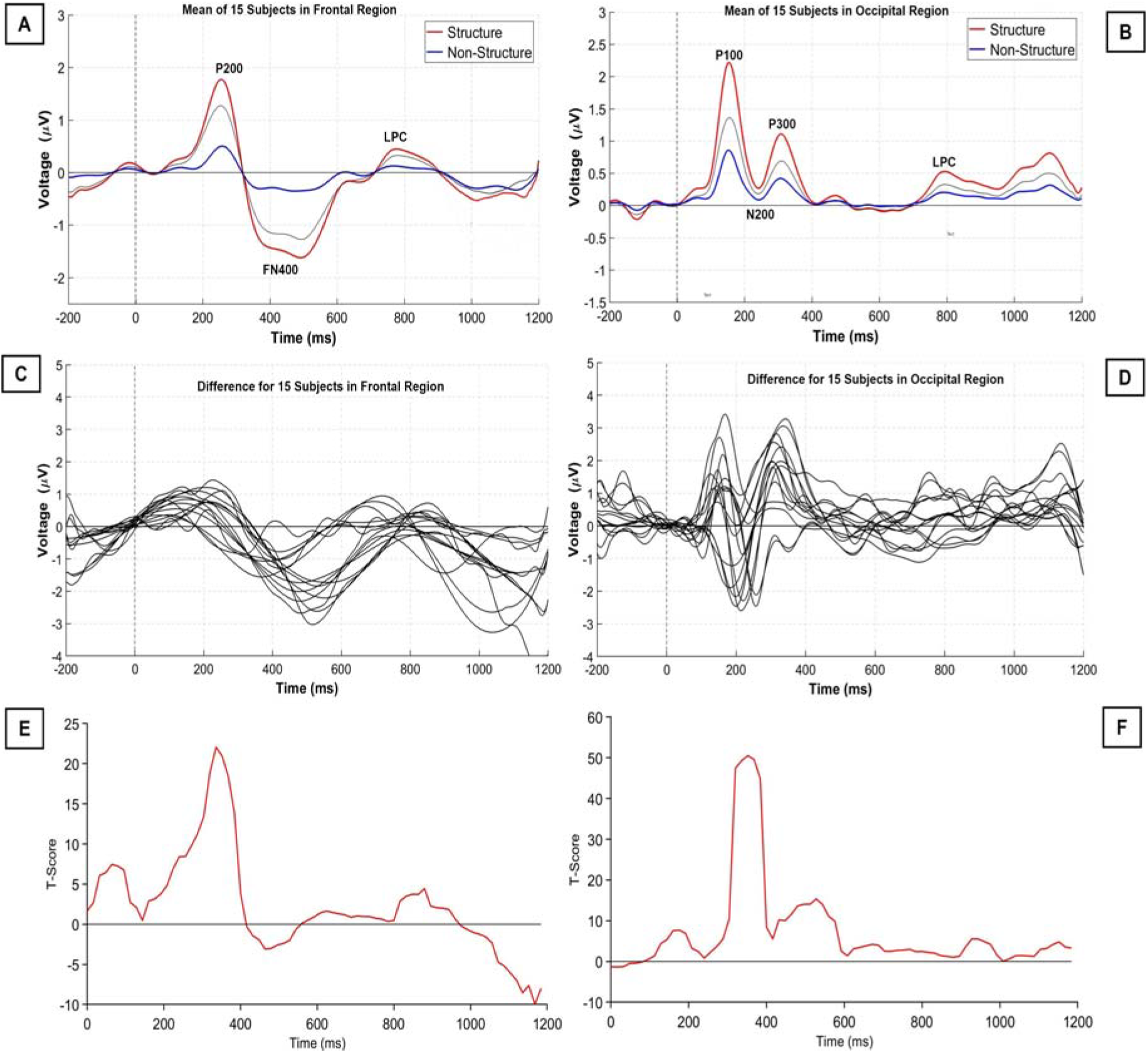
Grand ERP: (A) shows GERP for frontal region electrodes over 15 subjects, and (B) shows GERP for occipital region electrodes over 15 subjects (C) Difference curves averaged for all frontal electrodes for each of the 15 subjects (D) Difference curves averaged for all occipital electrodes for each of the 15 subjects (E) Mean T-Score for frontal electrodes over 15 subjects (F) T-Score for occipital electrodes over 15 subjects

The Grand ERP for Occipital region electrodes for 15 subjects is shown in Figure 6(B). The Statistical analysis [61]-[62] is done to evaluate the initial point of differential response across 15 subjects. The point of divergence for occipital region GERP occurred around 60 ms (electrode PO8), shown in Figure 6(B). All seven occipital electrodes showed consistent difference from 51ms (1.82 < t < 3.3). The difference curves for all 15 subjects is shown in Figure 6(D). The significance reaches a peak around 351ms where the mean t-score was 50.53 as illustrated in Figure 6(F).

### Differential response activation in primary visual cortex

It is believed that when the information enters through the visual sensory input, it is received by visual cortex area V1 [58]. That reception of information is confirmed by presence of C100 wave.

An ERP is calculated for all the electrodes in primary visual area to evaluate the processing of V1 area in the brain. The difference wave in Figure 5 showed first significant (*p<0*.*05*) negative peak around 110 ms after the stimulus onset which signifies the existence of C100 wave. The existence of C1 wave around 100 ms in Figure 5 confirms the reception of information by the visual primary cortex V1 region. Thus, it can be stated that V1 shows clear visual and statistical difference between the two classes.

### Information flow in brain

The flow of information can be judged by topographical maps presented in Figure 7. The topographical maps over the time-length of stimuli can be visualized and its representation in frontal and occipital ERP’s can be judged. The obtained point of divergence for two events in occipital and frontal regions is 60ms and 100 ms respectively which can also be stated by these brain activation maps. Thus, all the ERP processing can be visualized through these maps and the flow of information in the brain for perceptual grouping can be investigated.

**Figure 7.**
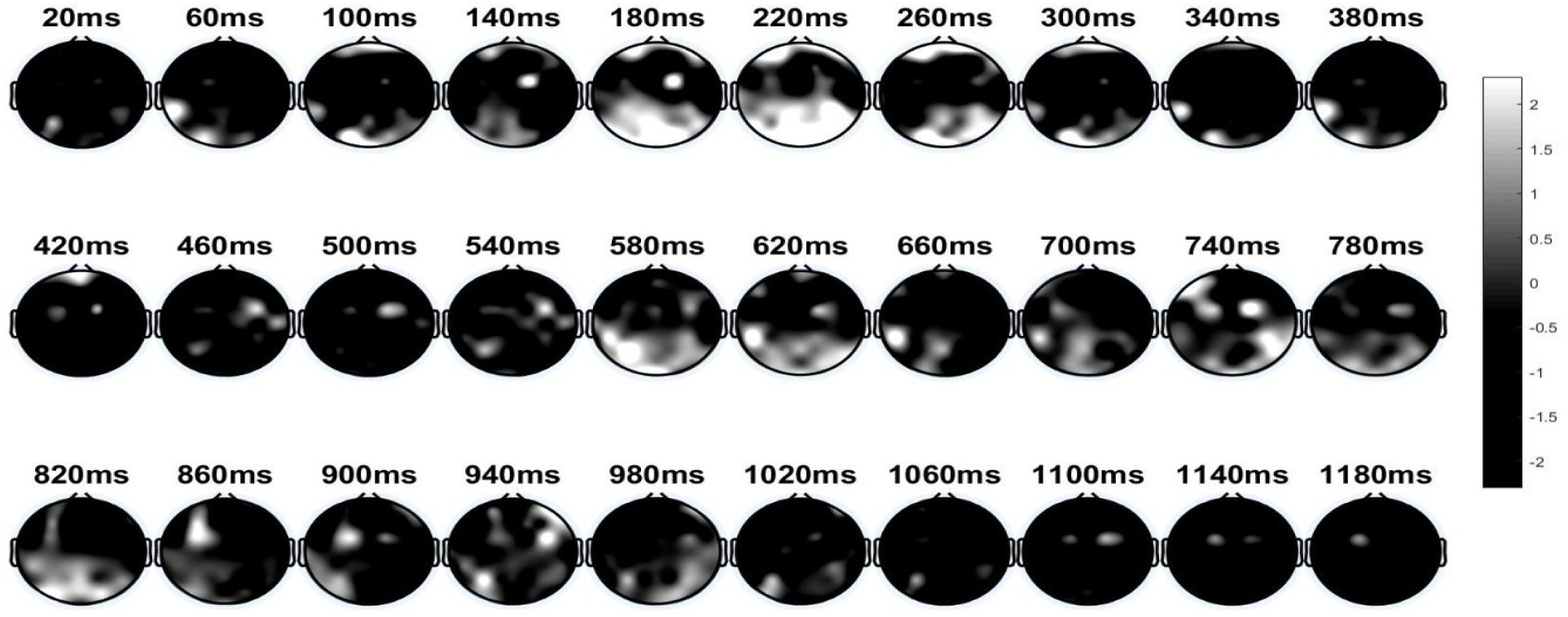
Topographical maps showing ERP difference between the events Structure and Non-Structure, where black color indicates no activity while white color represents the ERP activity.

The point of divergence in Occipital ERP in Figure 4(B) occurs around 60 ms which is reflected in topographical maps by an activation in occipital region around 60 ms. The reception of visual information may be the cause of this activation around 60 ms, which is confirmed by C100 peak in V1 region ERP [63] (in Figure 5). The point of divergence for Frontal ERP (Figure 5(A)) occurs around 100 ms, which is denoted by first activation in frontal region around 100 ms in topographical maps.

After 100 ms, an activation in occipital region can also be observed in topographical maps which is due to the difference in events for P100 wave in Occipital ERP. Thus, it indicates that the information is flowing from frontal region of brain to occipital region.

Around 180 ms, a high activation in frontal and occipital regions is obtained in topographical maps, which can be referred as frontal-occipital communication. This processing in the brain is also considered as the bottom-down processing of the brain.

### Relation between Synchrony and ERP divergence

PLV Synchrony for difference between the two events is shown in Figure 8. At 60 ms, an increase in connectivity in parieto-occipital region points to differential ERP divergence in occipital region. At 100 ms, increased connectivity in frontal region relates to point of divergence of differential ERP at frontal sites. Increased connectivity in occipital region around 140 ms, 220 ms, 340 ms and 800 ms is related to ERP difference peaks at occipital region. Enhanced connectivity in frontal region around 200 ms, 400 ms and 800 ms points to difference ERP peaks that occur at frontal region. Thus, it can be stated that the time-course of PLV relate to the ERP dynamics such that synchrony and ERP divergence occurs at the same time.

**Figure 8.**
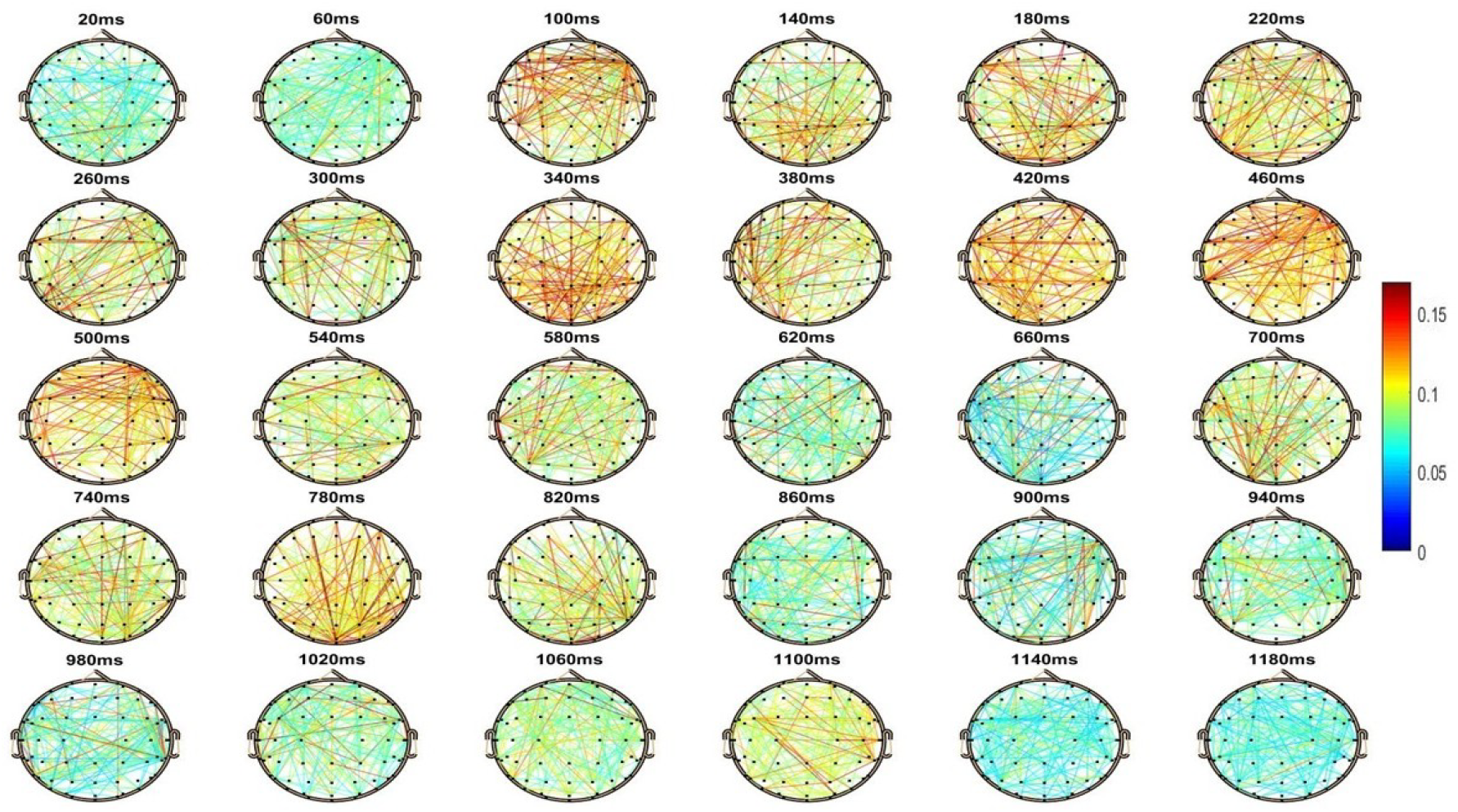
PLV Connectivity Maps for difference between the events over the time length of stimuli

## Discussion

Our results illustrates that there exist a visual as well as statistical difference between brain responses for structures and non-structures. It can be stated that perceptual grouping of dots as in case of structures produces high activation in brain as compared to the random dots. The straight lines composed of dot elements surrounded by noise composed of similar dots is detected such that the row of dots pops out the background [65]-[67], which reflects the figure-ground concept.

The obtained Frontal ERP signal constitutes of 3 components viz P200, N400, and a LPC (late positive component). The P200 component showed higher amplitude for grouping perception (i.e. Structures), shows more visual cognitive processing or more visual sensation for structures [68]-[69]. The higher amplitude N400 wave reflects potential for conceptual integration for structures [52]-[53]. The LPC elicits more attention in case of perceptual grouping as compared to random dots [54]. On the other hand, the obtained Occipital ERP signal constitutes of the following components viz P100, N200, P300, and a LPC (late positive component). The higher amplitude of P100 peak reflects a preliminary visual cognitive processing stage in the brain [55]. N200 component, responsible for cognitive control function represents higher amplitude peak for structures. The visual search during the figure-ground organization of line-segments amidst of random dots may be responsible for the higher values of N200 peak for structure trials [68]-[69]. The higher amplitude of P300 wave shows discrimination of one event from another [70]-[71]. Furthermore, the results showed the occurrence of N200 together with P300 which symbolizes the mental processes taking place to understand the random events [57]. Thus, the findings from ERP study extends the evidence of previous findings [22]. Also, the C1 wave in V1 region ERP showed that the primary visual cortex was involved at the early stage of grouping which obey previous findings as well [35]. The obtained behavioral reaction times dictates the maximum time required for visual processing of grouping perception while the ERP Analysis provides a stronger insight to the complete neural processing of perceptual grouping in the brain. Although, the mentioned findings provide deep insight about the processing in brain but does not clarify that how the different regions are connected during this processing. Thus, functional connectivity analysis [42]-[50] for difference between the events is performed. Therefore, this study can be considered as one study to investigate relation between synchrony time-course and ERP dynamics for perception grouping.

## Conclusion

In conclusion, the present study extends a deeper insight to existing perception grouping paradigm by manifesting electrophysiological and behavioral study to explore it. The neural processing and the biological markers related to grouping perception are evaluated and examined in frontal and occipital regions separately. ERP results suggest the perception of grouping induce higher activation in brain as compared to other stimuli. Also, the flow of information from V1 area to the frontal area of brain is found for structure class. Therefore, more specifically, it can be claimed that there exists a significant difference between the structure and non-structure evoked activations in brain. The C1 wave also showed the involvement of primary visual cortex in grouping perception. Also, the reaction time required by the brain for perceptual grouping in frontal and occipital regions is evaluated.

These findings obtained in this study are a basis of early visual processing and can be used as an additional feature for detection of neurological disorders. Furthermore, it would be interesting to examine how these findings emerge for neurophysiologically challenged subjects over longitudinal developmental trajectories.

